# The oldest peracarid crustacean reveals a Late Devonian freshwater colonisation by isopod relatives

**DOI:** 10.1101/2021.04.25.441336

**Authors:** N. Robin, P. Gueriau, J. Luque, D. Jarvis, A.C. Daley, R. Vonk

## Abstract

Peracarida (e.g., woodlice & side-swimmers) are, together with their sister-group Eucarida (e.g. krill & decapods), the most speciose group of modern crustaceans, suggested to have appeared as early as the Ordovician. While eucarids incursion onto land consists of mainly freshwater and littoral grounds, some peracarids have evolved fully terrestrial ground-crawling ecologies, inhabiting even our gardens in temperate regions (e.g. pillbugs and sowbugs). Their fossil record extends back to the Carboniferous and consists mainly of marine occurrences. Here, we provide a complete re-analysis of a fossil arthropod – *Oxyuropoda* – reported in 1908 from the Late Devonian floodplains of Ireland, and left with unresolved systematic affinities despite a century of attempts at identification. Known from a single specimen preserved in two-dimensions, we analysed its anatomy using digital microscopy and multispectral macro-imaging to enhance contrast of morphological structures. The new anatomical characters and completeness of *Oxyuropoda*, together with a phylogenetic analysis with representatives of all major Eumalacostraca groups, indicate that *Oxyuropoda* is a crown-peracarid, part of a clade including amphipods and isopods. As such, *Oxyuropoda* is the oldest known Peracarida, and provides evidence that derived peracarids had an incursion into freshwater and terrestrial environments as early as the Famennian, more than 360 million years ago.

## I. Introduction

Peracarid crustaceans (e.g., woodlice, opossum-shrimps, side-swimmers and comma-shrimps) are eumalacostracans that have diverged parallel to eucarids (shrimps, lobsters and crabs) to produce most modern crustacean diversity (67,000 described species) [1–3]. In peracarids, the most speciose groups are amphipods (side-swimmers; ∼10,000 species) and isopods (∼10,000 species), forming 20% of the diversity inhabiting rivers and lacustrine environments. A third of the isopod species are widespread terrestrial crawlers known as woodlice – common inhabitants of temperate gardens (e.g. pillbugs and sowbugs) [1–3]. While studies of more inclusive peracarid clades [4–9] did not suggest a diversification age for the group, that of the comparatively late-diverging peracarid order Isopoda is estimated by molecular analyses to have occurred during the mid to late Ordovician (∼455 Mya) [10,11], implying a long Palaeozoic history of Peracarida. However, reconciling the estimated Ordovician molecular time divergence of peracarids with their fossil record has been challenging [12,13], since the earliest fossil peracarids are of early to late Carboniferous age, consisting of a tanaid, [14,15], some putative mid-Carboniferous stem comma shrimps [16,17], and a late Carboniferous isopod [18,19].

A crustacean-looking arthropod, found in the early 20^th^ century from the Famennian (Late Devonian) of Ireland [20] has long been suspected of being an isopod-related animal. Reported from an undoubtedly freshwater assemblage, its affinities, if they could be determined, would help clarify the terrestrial evolution of Pancrustacea [20,21]. Its general aspect, reminiscent of the extant *Ligia*, a coastal terrestrial oniscoid woodlouse, earned it its name *Oxyuropoda ligioides* Carpenter and Swain, 1908. In addition to its round and short head fused to larger thoracic segments, it possesses a tapering and short pleon, with curved and pointed lateral pleurae on the abdominal segments, typical of isopods. However, its anatomy reveals six thoracic segments instead of the usual seven in isopods. Six such segments exist in other peracarids but these are very different in shape [22]. From its discovery to its last observation in 1985 [23] the fossil has successively been assigned to peracarids [20,24–29], phyllocarids [29], euthycarcinoids [30], arachnomorphs [31–33] or other arthropods [30–35]. Here, we re-analyze the visible and hidden anatomy of this continental (freshwater) arthropod using a combination of standard light photography, and newly developed luminescence-based imaging methods, and undertake a cladistic analysis to investigate its position within malacostracan crustaceans.

## 2. Material and methods

### (a) *Oxyuropoda* material and Kiltorcan Hill age

*Oxyuropoda ligioides* consists of a single specimen (part and counterpart; holotype NMING: F7633) housed in the palaeontological collection of the National Museum of Ireland, Dublin – Natural History, and recovered from Kiltorcan Hill. co. Kilkenny, Ireland. Besides *O. ligioides*, Kiltorcan Hill outcrops have yielded a range of continental freshwater-type organisms including lycophytes, progymnosperms, green algae *Bythotrephis*, freshwater bivalves, placoderms [20,36] and fragments of eurypterids [37–39]. Carpenter and Swain [20] reported *Oxyuropoda* from what can be recognized as the *Classic* or *Old quarry* of Kiltorcan [21,36,40]. The corresponding strata are encompassed by outcrops stratigraphically extending both below and above it, that do not yield macrofossils of the same quality and diversity, but miospore assemblages aiding in the age dating of the entire sequence. The most extensive outcrop to be found today on the hill is an active quarry referred as the *New quarry* [36] that yields LE miospore assemblages of the “Strunian” latest Famennian (uppermost Devonian). The topmost outcrop on the hill (*Roadstone quarry* [21,36,37]) yield a VI miospore assemblage of the lower most Carboniferous. The strata on Kiltorcan Hill therefore straddles the Devonian/Carboniferous Boundary [36] suggesting that the original Kiltorcan locality, located between New and Roadstone quarries horizons, is likely latest Famennian. To retrieve as many anatomical details as possible from this unique specimen, digital microscopy and multispectral macro-imaging have been carried out at the Institute of Earth Science, University of Lausanne, to enhance morphological contrast and reveal new anatomical features.

### (b) 2D and 3D digital microscopy

A Canon EOS 800D digital SLR camera fitted with an MP-E 65mm 1:28 1-5X Canon Macro Lens was used to photograph the specimen under artificial lighting from multiple directions. Polarising filters at the lens and the light source created crossed polarisation that reduced reflections and increased contrast. Digital microscopy images were collected using a Keyence VHX-7000 digital microscope equipped with a VH-ZOOT macro lens (0x to 50x magnification), connected to a VXH-7020 high performance 3.19-megapixel CMOS camera. High-resolution 2D images were collected through an automatic stitching process. 3D images presented herein in natural (**figure 1*e–g*, S1*c, e–g***) or false warm-cold (**figure S1*b***) colors were produced through an automatic vertical stacking process that creates 3D surface profiles.

**Figure 1.**
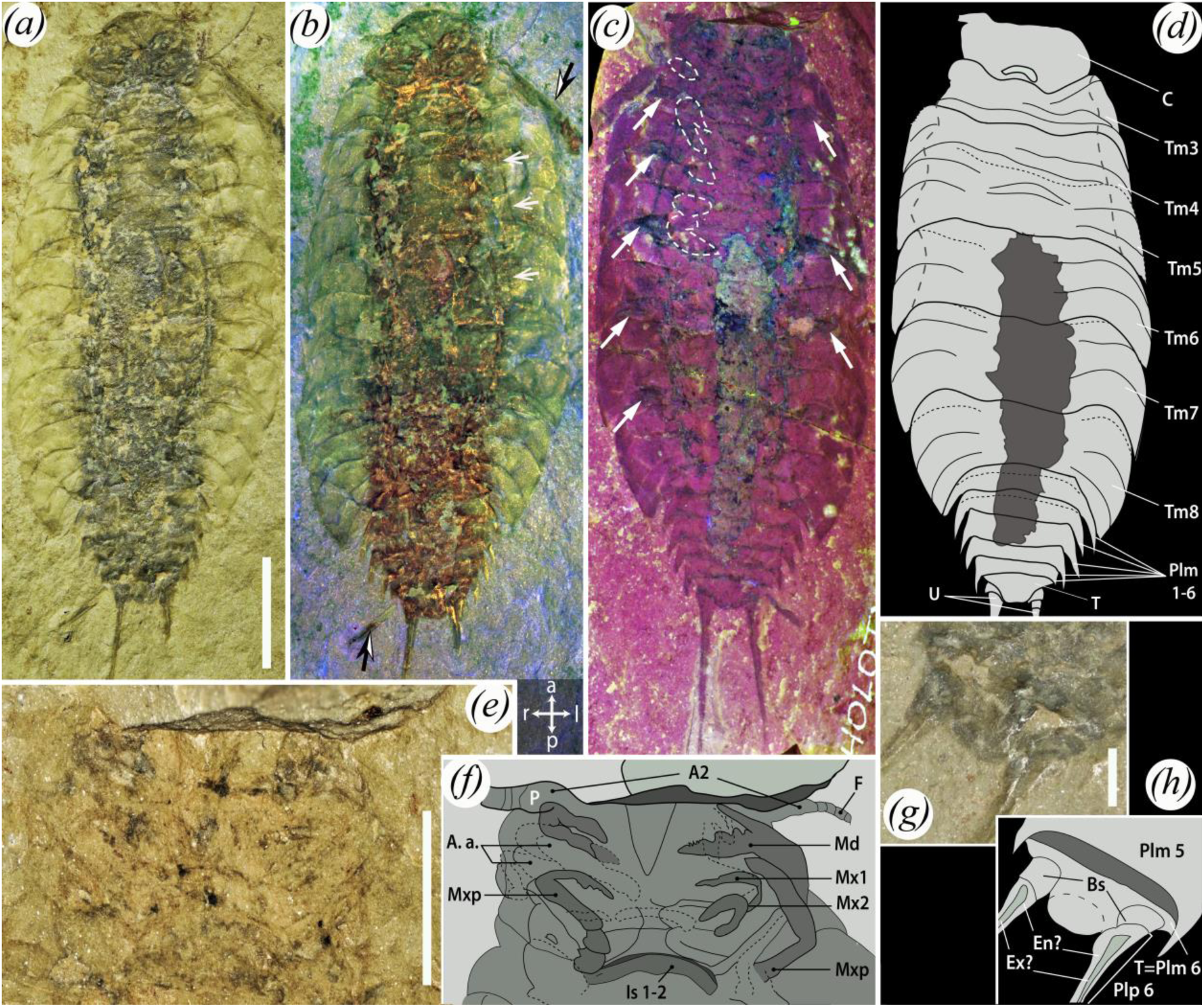
The anatomy of *Oxyuropoda ligioides* Carpenter and Swain, 1908, holotype NMING:F7633. Part (*a–b, g–h*, counterpart (*c–f*). **(*a–d*)**. Total body. **(*e–f*)**. Cephalon close-up. ***(g–h)***. Pleotelson close-up. Artificial lightning using crossed polarizing filters (*a*), Digital scanning surface microscopy (*e, g*) Multispectral macroimaging setting 1 (*b*), setting 2 (*c*), thin white dash-lines = outlines of the visible pereopods thin white dash-lines = outlines of the visible pereopods, long white arrows = areas of overlapping portions of thoracomeres, short white arrows = identified taphonomic cracks. Interpretative drawings (*f, h*), thin black dash-lines = limits from which pleurae are bending. **Captions**. A.a. Probable additionnal appendage, A2 = antenna, Bs = basipodites, C = cephalon, En? = possible endopodite, Ex? = possible exopodite, F = flagellum, Is 1–2 = intersternite 1–2 thickening, Md = possible mandible, Mxp = possible maxilliped, Mx1 = possible maxillula, Mx2 = possible maxilla, Tm 1–6 = thoracomere 1–6, P = antennal peduncle, Plm 1–6 = Pleomeres 1–6, Plp 6 = pleopod 6, T = telson, U = uropods. l/r/a/p = left, right, anterior, posterior referential. Scale bars = 10 mm (*a–d*), 5 mm, (*e–f*), 1 mm (g*–h*).

### (c) Multispectral Macroimaging

Building on the concept of UV photography [41] and visible light fluorescence imaging [42– 44], we collected reflection and luminescence images in various spectral ranges using an innovative imaging setup under development (see [45]). The setup consists of a low-noise 2.58-megapixel back illuminated sCMOS camera (PRIME 95B 25 mm, Photometrics) with high sensitivity from 200 to 1000 nm, fitted with a UV-VIS-IR 60 mm 1:4 Apo Macro lens (CoastalOptics) in front of which is positioned a filter wheel holding 8 Interference band-pass filters (Semrock) to collect images in 8 spectral ranges from 435 to 935 nm. Illumination is provided by 16 LED lights ranging from 365 up to 700 nm (CoolLED pE-4000), coupled to a liquid light-guide fiber fitted with a fiber-optic ring light-guide. As such, more than 90 different illumination/detection couples are available, and the resulting grayscale images can be combined into false-colour RGB images to enhance morphological contrasts or reveal new details in a wide range of fossils [45,46]. Stacking, image registration of the different couples, and production of false color RGB composites were performed using ImageJ. The field of view being smaller than the specimen, images of the full body were produced by producing RGB images at 3 different positions, which have then been stitched together using Image Composite Editor (Microsoft). False color RGB images presented herein were produced using two settings: (1) (**figure 1*b***) red – illumination 435 nm/detection 435 ±20 nm (reflection), green – illum. 660 nm/det. 650 ±30 nm (refl.), blue – illum. 435 nm/det. 571 ±36 nm (luminescence); (2) (**figure 1*c***) red – illum. 385 nm/det. 935 ±85 nm (lum.), green – illum. 435 nm/det. 571 ±36 nm (lum.), blue – illum. 435 nm/det. 435 ±20 nm (refl.).

### (d) Phylogenetic analysis

We investigated the phylogenetic position of †*Oxyuropoda* using a morphological dataset for malacostracan crustaceans modified from [5]. The phylogenetic significance of uropods in eumalacostracans was recently evidenced in [47]. Consequently, we scored four extra uropodal characters that are preserved in *Oxyuropoda*, following the approach of [48] (see SI). Likewise, because amphionidacean crustaceans were recently demonstrated to be decapod larvae instead of proper distinct taxa [12,48,49], the operational taxonomic unit (OTU) *Amphionidacea* was culled from our analyses. Final data matrices after [5] (25 OTUs with 181 adult morphological characters) were built in Mesquite 3.51 [50]. We included in a second analysis †*Tealliocaris*, a Late Devonian-Carboniferous eumalacostracan with debated peracarid or decapod affinities [12,51–56] that has been reported from late Fammenian (VCo Oppel biozone) freshwater (or at least continental water) horizons of Belgium [57,58]. Undetermined and not preserved characters were scored as ‘?’, and inapplicable characters as ‘–‘. Multiple character states present in a given OTU were scored as polymorphisms. We analysed the data set using Bayesian Inference as implemented in MrBayes v.3.2.6 [59]. The dataset was analysed under the traditional Mk model [60] with an ascertainment bias correction to account for scoring only variable morphological characters, and gamma distributed rate variation. Each analysis was performed with two independent runs of 3×10^7^ generations each. We used the default settings of four chains (one cold and three heated) per each independent run. The relative burn-in fraction was set to 25% and the chains were sampled every 200 generations. Temperature parameter was set to 0.01 as determined by preliminary runs to achieve chain mixing values in the optimal range (0.4 – 0.8). Convergence of independent runs was assessed through the average standard deviation of split frequencies (ASDSF << 0.01) and potential scale reduction factors [PSRF ≈ 1 for all parameters [61]]. We used Tracer v.1.7.1 [62] to determine whether the runs reached stationary phase and to ensure that the effective sample size (ESS) for each parameter was greater than 200. Results of the Bayesian runs were summarized as a majority-rule consensus tree of the post-burnin sample (**figure 2, S2**). The obtained tree branches were constrained over geological time in our figures respectively to estimated divergence age obtained or recommended from morphological and molecular studies on Branchiopoda [63], Isopoda [10,11,13] and Amphipoda [64].

**Figure 2.**
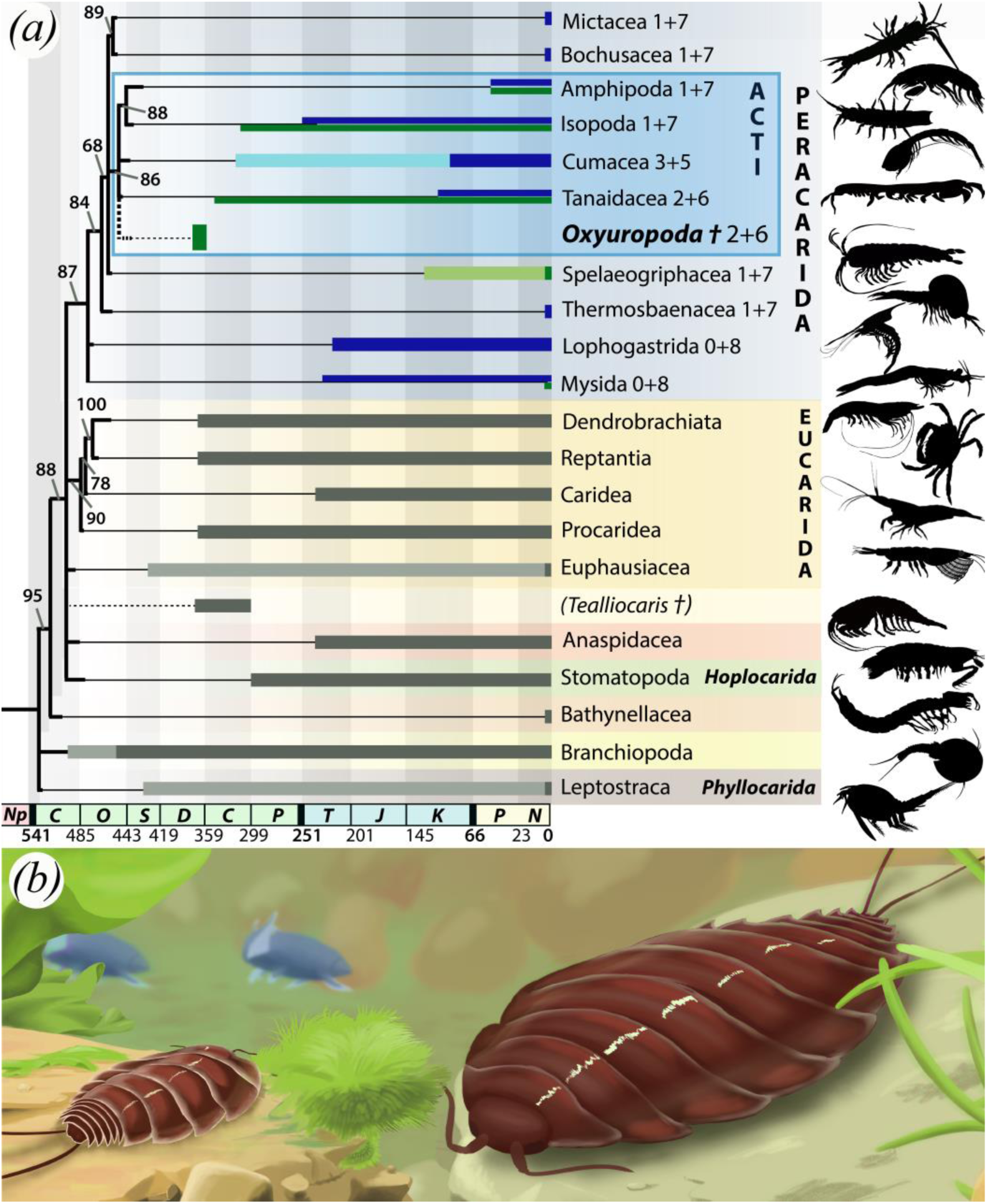
Phylogeny of the Malacostraca including *Oxyuropoda ligioides* Carpenter and Swain, 1908 after [1] and reconstruction of the animal in its freshwater environment in the Late Devonian. **(*a*)**. Bayesian majority-rule consensus topology and branch lengths of the post-burnin sample of trees, plotted on geological times. Branches with posterior probability support ≤60% not shown. The obtained tree (black) accommodates here: (1) the Late Ordovician divergence of Isopoda, (2) the minimal Late Carboniferous divergence of Amphipoda, (3) the Early Cambrian clade age of Branchiopoda. Light and dark thick lines respectively for stem - and crown-groups; grey by default, blue for peracarid marine taxa, green for peracarid freshwater taxa, see [2] for age justification). ACTI = clade grouping Amphipoda, Cumacea, Tanaidacea and Isopoda. 0+8: code for *number of malacostracan thoracomeres integrated in the cephalon* + *number of thoracomeres in the thorax*. (see S2*a* for detail). Added topological location of *Tealliocaris* when included in the analysis (see S2*b* for detail). **(*b*)**. Amended reconstruction of two *Oxyuropoda ligioides* in the Kiltorcan Old Quarry floodplains of the Upper Famennian (Upper Devonian), Kilkenny co., Leinster, Ireland. In association with the onland progymnosperms *Archeopteris hibernica*, and underwater algae *Bythotrephis* sp., the placoderms *Glyptolepis leptopterus*, and freshwater bivalve *Archanodon jukesi*. Reconstruction by Diane Dabir Moghaddam.

### 3. Results and discussion

The advanced imaging of the only known *Oxyuropoda ligioides* specimen resolved both new anatomical features, never observed by optical microscopy, and the artifacts leading to previous erroneous interpretations of the fossil (**figure 1*a–h***). Elongated elements have earlier been questioned to correspond to either to antennae/uropod parts or to plant remains that look very similar to the animal appendages and are very abundant in the specimen [20,32]. Some of these are now surely assigned to plants observed detached from, and underneath, the arthropod body (black and white arrows in **figure 1*b***). A 3D surface rendering using digital microscopy accentuated a different extent of deformation between the right and left pleurae, with the left ones having bent more vertically than the right ones (**figure S1*b***). Parallel to the left pleurae, the specimen exhibits a marked groove that we interpret as a break associated with the most pronounced compression of the left side of the thorax (**figure 1*b***). This localised bending implies that the body of *Oxyuropoda* was generally flattened dorsoventrally, but had the extremities of its pleurae directed ventrally rather than laterally. Elements of the cephalon previously suggested to be the eyes are here revealed to be topographical artifacts (thick dash lines in **figure S1*e***), that we cannot exclude to reflect more ventrally located organs, but that we rather interpret as local, and asymmetric, bulges reflecting superposition of appendages. The first segments of the antennae – with large peduncles (**figure 1*e–f***) – and the outlines of the buccal appendages are observed for the first time (**figure 1*e–f*, S1*c–f***). These are comparable to the derived appendages of modern malacostraca, being elongated in shape (*pediform* following [65]) and geniculated, which we interpret as maxillipeds, maxillae (mx 1 and mx 2) and mandibles, as well as a probable pair of additional appendages that hard to define in outline (**figures 1>*e–f***). The mandible structure shows a strong incisor, crenelated at its very tip end and bearing 4 teeth, as well as an elongated palp directed forward (**figure 1*e–f***). The number of these appendages suggests that there are three somites forming the cephalon part of the cephalothorax (despite rather reduced cephalon dimensions, so likely thoracomeres 1 and 2 are fused, **figure 1*a–d***); the presence/absence of a carapace above these segments cannot be established. The thorax comprises six thoracomeres (highlighted by short regions of overlapping proximal pleurae, **figure 1c**) increasing in size from anterior to posterior (figure 1*a–d*, S1*a*), each crossed by two transverse suture ridges and likely bearing a pair of pereopods which basis are locally visible (**figure 1*c*, S1*c–f***). Some pleurae exhibit longitudinal to oblique grooves, evoking the coxal articulations found in Isopoda thoracomeres. The close examination of these structures reveal they correspond to be taphonomic cracks, meaning there is no anatomical evidence for coxal plate articulations in *Oxyuropoda* (**figure S1g**). The pleon is composed of six pleomeres (**figure 1*a–d*, S1a**), with the last one bearing the telson and a pair of uropods posteriorly oriented and styliform in shape (**Figure 1*g–h***). See ESM for systematic palaeontology and full description, including measurements.

A cephalothorax, six pleonal segments (excluding telson) and uropods are malacostracan features [5,7,8,65–68], proper to all lineages but Leptostraca, while a stout tooth-like mandibular incisor is not found in these nor within Bathynellacea [8]. The absence of furcal extensions on the telson is also improper to Hoplocarida; these being indeed displayed in some extinct representatives [8]. The incorporation of a number of anterior thoracomeres into the cephalon is typical of all other malacostracan clades [5,7,8]. The body plan of *Oxyuropoda* displays only seven directly observable cephalothoracic somites out of the nine typical of malacostracans, requiring further digging into other diagnostic characters such as its uropodal features [47]. In eumalacostracan crustaceans, uropods represent a modification of the sixth pair of pleonal appendages, which in contrast to caudal furca, are real, articulated appendages with a basipodite; they are posteriorly oriented and styliform in shape in *Oxyuropoda*. The basipodite is straight without extensions, and the uropod without carinae (contrary to interpretation in [25], ridge corresponding to exo/-endopodites division), a combination of features found in Bathynellacea and in lately diverged peracarids (Spelaeogriphacea, Mictacea, Cumacea, Tanaidacea, Isopoda and Amphipoda) [8,47].

Bayesian inference phylogenetic analysis places *Oxyuropoda* among a strongly supported late diverged clade of Peracarida (posterior probability 86) – previously found in [6,7,69] – that includes Amphipoda, Cumacea, Tanaidacea, and Isopoda (referred as clade ACTI; **figures 2*a*, S2*a***). The internal phylogenetic relationships of these main peracarid clades, with the exception of a few more often recovered patterns (e.g. Mysida grouped with Lophogastrida), are so far not resolved [4–9,13,65–67,69–74]. The input of *Oxyuropoda* maintains (1) the sister-relationship of Isopoda and Amphipoda recovered in [6–8,65,67,69,73,74] as well as (2) the earliest divergence of the marine Mysidacea (Mysida and Lophogastrida) [5–8,65,67,69,71– 74], followed by those of (3) marine Thermosbaenacea and (4) a mixed marine and continental clade comprising Spelaeogriphacea, ACTI and clade Mictacea/Bochusacea. *Tealliocaris*, whose affinities remain unresolved in our analysis, is found outside of modern peracarid clades **(figures 2*a*, S2*b***). With its late Devonian age, *Oxyuropoda* is the oldest known crown-group Peracarida, outdating *Anthracocaris scotica* Peach, 1882 from the Visean (Early Carboniferous) of the Calciferous Sandstone of the UK [14], which has been identified as a crown-Tanaidacea (Anthracocarididae, see [15]). No stem-peracarid lineage has so far been identified [75]. Based on the information currently available, the molecular/fossil-based supported diversification time of amphipods & isopods on one side [10,11,64], and of branchiopods on the other [63], suggest a first peracarid diversification in the latest Cambrian (**figure 2*a***). Recovering *Oxyuropoda* (2 thoracomeres integrated in the cephalon and 6 in the thorax) as a crown peracarid may help polarize the direction of change of cephalic integration of anterior thoracomeres among Peracarida, yet this remains tentative until intrarelationships within crown Peracarida are better resolved.

Finally, the affini ty of *Oxyruropoda* gives insight into the timing of colonization of non-marine environments by peracarids. *Oxyuropoda* plots either as the direct sister taxon, or within a sister clade (clade ACTI for Spelaeogriphacea) of groups with partial or strict freshwater representatives, the record of which even extends to their fossil representatives (e.g. Tanaidacea, Isopoda, Spelaeogriphacea) (**figure 2*a***). Being likely part of an independent lineage of peracarids, *Oxyuropoda* provides further evidence that derived Peracarida were present in continental settings as early as the Fammenian, implying the fast colonisation of continental waters (and land?) during the evolution of clade ACTI and likely of Spelaeogriphacea. Overall, our results indicate that besides Branchiopoda ([57,76–80], and regardless of the affinities of tealliocaridids, highly derived vericrustacean groups had already colonized continental ecosystems by the Late Devonian (Famennian, more than 360 Ma) (**figures 2*a–b*, S3**).

## ADDITIONNAL INFORMATION

### DATA ACCESSIBILITY

Requests for access to the fossil material should be addressed to Nigel Monaghan of the National Museum of Ireland, Natural History, Dublin. Additional data are available as electronic supplementary material.

### AUTHORS’ CONTRIBUTIONS

N.R. conceived the project and applied for its funding. P.G. A.C.D. and N.R. performed digital microscopy and multispectral microimaging, R.V. and N.R. described the fossil anatomy, N.R. and P.G. defined the phylogenetic characters and coded the matrix. J.L., conducted the phylogenetic analyses, E.J. provided the geological and age context of the fossil. N.R. drafted the manuscript with the contributions of other authors.

### FUNDING

This study was supported by a Sylvester-Bradley Award 2020-PA-SB201902: Investigating the enigmatic Devonian arthropod *Oxyuropoda* — of the Palaeontological Association. N.R. fellowship at UCC is supported by a Government of Ireland Post-doctoral Fellowship of the Irish Research Council-MG21562. The multispectral microimaging system used herein was funded by the Swiss National Science Foundation under grant number 205321_179084 “Arthropod Evolution during the Ordovician Radiation: Insights from the Fezouata Biota”, awarded to A.C.D.

## ACKNOWLEDGEMENTS

The authors warmly thank the palaeoartist D. Dabir-Moghaddam who produced the first scientific reconstruction of the animal. We thank A. Bardon for thoughtful peracarid talks, and F. Noailles for bibliographic support. N.R wishes to thank the ANOM lab members for their kind pre-COVID hosting, as well as M. McNamara for her immediate support in this somehow different project. N.R. thanks the Palaeontological Association for accepting to fund this initiative. We are particularly grateful to palaeontologist and curator M. Parkes for enabling the study of this material and very kindly allowing each aspect of its physical analysis. M. Parkes passed away in 2020; we dedicate this study to him.

## References

1. Wilson GDF. 2007 Global diversity of Isopod crustaceans (Crustacea; Isopoda) in freshwater. In Freshwater animal diversity assessment, pp. 231–240. Springer.

2. Väinölä R, Witt JDS, Grabowski M, Bradbury JH, Jazdzewski K, Sket B. 2007 Global diversity of amphipods (Amphipoda; Crustacea) in freshwater. In Freshwater Animal Diversity Assessment, pp. 241–255. Springer.

3. Arfianti T, Wilson S, Costello MJ. 2018 Progress in the discovery of amphipod crustaceans. PeerJ 2018, 1–16. (doi:10.7717/peerj.5187)

4. Watling L. 1999 Toward understanding the relationships of the peracaridan orders: the necessity of determining exact homologies. In Crustaceans and the Biodiversity Crisis: Proceedings of the Fourth International Crustacean Congress, 1999, pp. 73–89.

5. Richter G, Scholtz. 2001 Phylogenetic analysis of the Malacostraca (Crustacea). J. Zool. Syst. Evol. Res. 39, 113–136. (doi:10.1046/j.1439-0469.2001.00164.x)

6. Poore GCB. 2005 Peracarida: monophyly, relationships and evolutionary success. Nauplius 13, 1–27.

7. Jenner RA, Dhubhghaill CN, Ferla MP, Wills MA. 2009 Eumalacostracan phylogeny and total evidence: limitations of the usual suspects. BMC Evol. Biol. 9, 1–20.

8. Wills MA, Jenner RA, Dhubhghaill CN. 2009 Eumalacostracan evolution: conflict between three sources of data. Arthropod Syst. Phylogeny 67, 71–90.

9. Wilson G. 2009 The phylogenetic position of the Isopoda in the Peracarida (Crustacea: Malacostraca). Arthropod Syst. Phylogeny 67, 159–198.

10. Lins LSF, Ho SYW, Wilson GDF, Lo N. 2012 Evidence for Permo-Triassic colonization of the deep sea by isopods. Biol. Lett. 8, 979–982. (doi:10.1098/rsbl.2012.0774)

11. Lins LSF, Ho SYW, Lo N. 2017 An evolutionary timescale for terrestrial isopods and a lack of molecular support for the monophyly of Oniscidea (Crustacea: Isopoda). Org. Divers. Evol. 17, 813–820. (doi:10.1007/s13127-017-0346-2)

12. Hegna TA, Luque J, Wolfe JM. 2020 The fossil record of the Pancrustacea. In The natural History of Crustacea - Volume 8 Evolution and biogeography (eds M Thiel, G Poore), pp. 21–52. Oxford University Press.

13. Broly P, Deville P, Maillet S. 2013 The origin of terrestrial isopods (Crustacea: Isopoda: Oniscidea). Evol. Ecol. 27, 461–476. (doi:10.1007/s10682-012-9625-8)

14. Calman WT. 1933 On Anthracocaris scotica (Peach), a fossil Crustacean from the Lower Carboniferous. Ann. Mag. Nat. Hist. 11, 562–565. (doi:10.1080/00222933308673688)

15. Schram FR, Sieg J, Malzan E. 1986 Fossil Tanaidacea. Trans. San Diego Soc. Nat. Hist. 21, 127– 144.

16. Schram FR. 2003 Paleozoic cumaceans (Crustacea, Malacostraca, Peracarida) from North America. Contrib. to Zool. 72, 1–16.

17. Luque J, Gerken S. 2019 Exceptional preservation of comma shrimp from a mid-Cretaceous Lagerstätte of Colombia, and the origins of crown Cumacea. Proc. R. Soc. B Biol. Sci. 286. (doi:10.1098/rspb.2019.1863)

18. Schram FR. 1970 Isopod from the Pennsylvanian of Illinois. Science (80-.). 169, 854–855.

19. Schram FR. 1974 Paleozoic Peracarida of North America. Fieldiana Geol. 33, 1–95.

20. Carpenter GH, Swain I. 1908 A new Devonian isopod from Kiltorcan, County Kilkenny. In Proceedings of the Royal Irish Academy. Section B: Biological, Geological, and Chemical Science, pp. 61–67.

21. Jarvis DE. 2000 Palaeoenvironment of the plant bearing horizons of the Kiltorcan Hill, Co. Kilkenny, Ireland. New Perspect. Old Red Sandstone, 8719.

22. Thiel M, Hinojosa I. 2009 Peracarida – Amphipods, Isopods, Tanaidaceans & Cumaceans. Mar. Benthic Fauna Chil. Patagon., 671–718.

23. Almond Lawson JD. 1985 The Silurian-Devonian fossil record of the Myriapoda. Philos. Trans. R. Soc. London. B, Biol. Sci. 309, 227–237.

24. Schmidt C, Leistikow A. 2004 Catalogue of genera of the terrestrial Isopoda (Crustacea: Isopoda: Oniscidea). Steenstrupia 28, 1–118. (doi:10.1177/004728758202000301)

25. Rolfe WDI. 1969 Arthropoda incertae sedis. In Treatise on invertebrate paleontology part R. (ed RC Moore), pp. 620–625. Washington: Geological Society of America.

26. Vandel A, de Barros Machado A. 1946 Crustacés isopodes terrestres (Oniscoïdea) épigés et cavernicoles du Portugal.

27. Reiff E. 1936 Isopoden aus dem Lias Delta (Amaltheenschichten) Schwabens. Palaeontol. Zeitschrift 18, 49–90. (doi:10.1007/BF03041710)

28. Calman WT. 1909 IV. On the Anaspidacea, living and fossil. By Geoffrey Smith. Quarterly Journal of Microscopical Science, vol. liii, pt. iii, May, 1909. Geol. Mag. 6, 425–426.

29. Roger J. 1953 Sous-classe des Malacostraces (Malacostraca Latreille, 1806). In Traite de Paleontologie (ed J Piveteau), pp. 303–378. Elsevier Masson SAS.

30. Schram FR. 1971 A strange arthropod from the Mazon Creek of illinois and the trans Permo-triassic Merostomoidea (Trilobitoidea). Fieldiana Geol. 20, 85–102.

31. Broili F. 1932 Eine neue Crustacee aus dem rheinischen Unterdevon. Sitzungsb. d. math.-naturw. Abt. 1, 27–38.

32. Schultze P. 1939 Bemerkenswerte palaeozoische arthropoden, die wahrscheinlich in die spinnentierreihe gehören. Zoomorphology 35, 169–182.

33. Størmer L. 1944 On the relationships and phylogeny of fossil and recent Arachnomorpha. A comparative study on Arachnida, Xiphosura, Eurypterida, Trilobita, and other fossil Arthropoda. Skr. Nor. Vidensk. Acad. Oslo, L Mat.-Naturvidensk. Klasse 1944 I5, 1–158.

34. McCoy VE, Strother PK, Briggs DEG. 2012 A possible tracemaker for Arthrophycus alleghaniensis. J. Paleontol. 86, 996–1001. (doi:10.1666/11-133r1.1)

35. McNamara KJ, Trewin NH. 2002 A Euthycarcinoid arthropod from the Silurian of western Australia. Palaeontology. 36, 319–335.

36. Jarvis E. 1990 New palynological data on the age of the Kiltorcan Flora of Co. Kilkenny, Ireland. J. Micropalaeontology 9, 87–94. (doi:10.1144/jm.9.1.87)

37. Clayton G, Graham JR, Higgs K, Holland CH, Naylor D. 1979 Devonian rocks in Ireland: a review. J. Earth Sci., 161–183.

38. Hellier Baily W. 1877 On Fossils from the Upper Old Red Sandstone of Kiltorcan Hill, in the County of Kilkenny. Report No. 1. Proc. R. Irish Acad. Sci. 2, 45–48.

39. Cole GAJ. 1901 II.—On Belinurus kiltorkensis, Baily. Geol. Mag. 8, 52–54.

40. Colthurst JRJ, Jrj C. 1978 The geology of the Lower Palaeozoic and Old Red Sandstone rocks of the Slievenamon Inlier, Counties Tipperary and Kilkenny.

41. Tischlinger H. 2001 Die oberjurassischen Plattenkalke von Daiting. Klass. Fundstellen Der Palaontologie, Bd. IV. Korb Goldschneck Verlag, 139–151.

42. Haug C, Haug JT, Waloszek D, Maas A, Frattigiani R, Liebau S. 2009 New methods to document fossils from lithographic limestones of southern Germany and Lebanon. Palaeontol. Electron. 12.

43. Haug JT et al. 2011 Autofluorescence imaging, an excellent tool for comparative morphology. J. Microsc. 244, 259–272.

44. Kaye TG, Falk AR, Pittman M, Sereno PC, Martin LD, Burnham DA, Gong E, Xu X, Wang Y. 2015 Laser-stimulated fluorescence in paleontology. PLoS One 10, e0125923.

45. Brayard A, Guériau P, Thoury M, Escarguel G. 2019 Glow in the dark: Use of synchrotron μXRF trace elemental mapping and multispectral macro-imaging on fossils from the Paris Biota (Bear Lake County, Idaho, USA). Geobios 54, 71–79.

46. Klug C, Landman NH, Fuchs D, Mapes RH, Pohle A, Guériau P, Reguer S, Hoffmann R. 2019 Anatomy and evolution of the first Coleoidea in the Carboniferous. Commun. Biol. 2, 1–12.

47. Kutschera V, Maas A, Waloszek D. 2012 Uropods of eumalacostraca (Crustacea s.l.: Malacostraca) and their phylogenetic significance. Arthropod Syst. Phylogeny 70, 181–206.

48. Gueriau P, Rak S, Broda K, Kumpan TT. V, Valach P, Zatoń M, Charbonnier S, Luque J. 2021 Exceptional Late Devonian arthropods document the origin of decapod crustaceans. Biol. Lett.

49. De Grave S, Chan T-Y, Chu KH, Yang C-H, Landeira JM. 2015 Phylogenetics reveals the crustacean order Amphionidacea to be larval shrimps (Decapoda: Caridea). Sci. Rep. 5, 1–8. (doi:10.1038/srep17464)

50. Maddison WP. 2018 Mesquite: a modular system for evolutionary analysis. Evolution (N. Y). 62, 1103–1118.

51. Briggs DEG, Clarkson ENK. 1985 The Lower Carboniferous shrimp Tealliocaris from Gullane, East Lothian, Scotland. Trans. R. Soc. Edinb. Earth Sci. 76, 173–201. (doi:10.1017/S0263593300010439)

52. Taylor RS, Yan-Bin S, Schram FR. 1998 New pygocephalomorph crustaceans f rom the Permian of China and their phylogenetic relationships. Palaeontology 41, 815–834.

53. Clark N. 2013 Tealliocaris: a decapod crustacean from the Carboniferous of Scotland. Palaeodiversity 6, 107–133.

54. Gueriau P, Charbonnier S, Clément G. 2014 First decapod crustaceans in a Late Devonian continental ecosystem. Palaeontology 57, 1203–1213. (doi:10.1111/pala.12111)

55. Jones WT, Feldmann RM, Schram FR, Schweitzer CE, Maguire EP. 2016 The Proof is in the Pouch: Tealliocaris is a Peracarid. Palaeodiversity 9, 75–88. (doi:10.18476/pale.v9.a5)

56. Yang Q, Gueriau P, Charbonnier S, Ren D, Béthoux O. 2018 A new tealliocaridid crustacean from the Late Carboniferous of North China and its biogeographic implications. Acta Palaeontol. Pol. 63.

57. Gueriau P, Rabet N, Olive S, Be O, Lagebro L, Vannier J, Briggs DEG. 2016 A 365-million-year-old freshwater community reveals morphological and ecological stasis in branchiopod crustaceans. Curr. Biol. 26, 383–390. (doi:10.1016/j.cub.2015.12.039)

58. Denayer J, Prestianni C, Gueriau P, Olive S, Clement G. 2016 Stratigraphy and depositional environments of the Late Famennian (Late Devonian) of Southern Belgium and characterization of the Strud locality. Geol. Mag. 153, 112–127.

59. Ronquist F et al. 2012 MrBayes 3.2: efficient Bayesian phylogenetic inference and model choice across a large model space. Syst. Biol. 61, 539–542.

60. Lewis PO. 2001 A likelihood approach to estimating phylogeny from discrete morphological character data. Syst. Biol. 50, 913–925.

61. Gelman A, Rubin DB. 1992 Inference from iterative simulation using multiple sequences. Stat. Sci. 7, 457–472.

62. Rambaut A, Drummond AJ, Xie D, Baele G, Suchard MA. 2018 Posterior summarization in Bayesian phylogenetics using Tracer 1.7. Syst. Biol. 67, 901.

63. Wolfe JM, Daley AC, Legg DA, Edgecombe GD. 2016 Fossil calibrations for the arthropod Tree of Life. Earth-Science Rev. 160, 43–110. (doi:10.1016/j.earscirev.2016.06.008)

64. Copilaş-Ciocianu D, Borko Š, Fišer C. 2020 The late blooming amphipods: Global change promoted post-Jurassic ecological radiation despite Palaeozoic origin. Mol. Phylogenet. Evol. 143, 106664. (doi:10.1016/j.ympev.2019.106664)

65. Schram FR. 1984 Relationships Within Eumalacostracan Crustacea. Trans. San Diego Soc. Nat. Hist. 20, 301–312. (doi:10.5962/bhl.part.29008)

66. Dahl E. 1983 Malacostracan phylogeny and evolution. Crustac. Phylogeny 1, 189–212.

67. Wills MA. 1998 A phylogeny of recent and fossil Crustacea derived from morphological characters. In Arthropod Relationships, pp. 189–209.

68. Dahl E. 1992 Aspects of Malacostracan Evolution. Acta Zool. 73, 339–346. (doi:10.1111/j.1463-6395.1992.tb01104.x)

69. Wagner HP. 1994 A monographic review of the Thermosbaenacea (Crustacea: Peracarida). Zool. Verh. Leiden 291, 1–338.

70. Siewing R. 1963 Studies in malacostracan morphology: results and problems. In Phylogeny and evolution of Crustacea, pp. 85–103. Museum Comp. Zool., Harvard Univ.

71. Watling L. 1981 An Alternative Phylogeny of Peracarid Crustaceans. J. Crust Biol. 1, 201–210. (doi:10.2307/1548159)

72. Wheeler WC. 1998 Sampling, groundplans, total evidence and the systematics of arthropods. In Arthropod relationships, pp. 87–96. Springer.

73. Schram FR, Hof CHJ. 1998 Fossils and the interrelationships of major crustacean groups. In Arthropod Fossils and Phylogeny (ed GD Edgecombe), p. 233. Cambridge University Press.

74. Watling L, Hof CHJ, Schram FR. 2000 The place of the Hoplocarida in the Malacostracan Pantheon. J. Crustac. Biol. Biol. 20, 1–11. (doi:10.1163/1937240X-90000002)

75. Vicente CS, Cartanyà J. 2017 A new mysid (Crustacea, Mysida) from the Ladinian Stage (Middle Triassic) of Conca de Barberà (Catalonia, NE Iberian Peninsula). J. Paleontol. 91, 968–980. (doi:10.1017/jpa.2017.24)

76. Walossek D. 1993 The upper Cambrian Rehbachiella and the phylogeny of Branchiopoda and Crustacea. Lethaia 26, 318.

77. Anderson LI, Crighton WRB, Hass H. 2003 A new univalve crustacean from the Early Devonian Rhynie chert hot-spring complex. Earth Environ. Sci. Trans. R. Soc. Edinburgh 94, 355–369.

78. Fayers SR, Trewin NH. 2002 A new crustacean from the Early Devonian Rhynie chert, Aberdeenshire, Scotland. Earth Environ. Sci. Trans. R. Soc. Edinburgh 93, 355–382.

79. Scourfield DJ. 1926 V. On a new type of crustacean from the old red sandstone (Rhymie Chert Bed, Aberdeenshire)—Lepidocaris rhyniensis, gen. et sp. nov. Philos. Trans. R. Soc. London. Ser. B, Contain. Pap. a Biol. Character 214, 153–187.

80. Trewin NH, Fayers SR, Kelman R. 2003 Subaqueous silicification of the contents of small ponds in an Early Devonian hot-spring complex, Rhynie, Scotland. Can. J. Earth Sci. 40, 1697–1712.

